# Comparative Analysis of Structural and Dynamical Properties of Lipid Membranes Simulated with the AMBER Lipid21 Force Field Using SPC/E, TIP3P, TIP3P-FB, TIP4P-FB, TIP4P-Ew, TIP4P/2005, TIP4P-D, and OPC Water Models

**DOI:** 10.64898/2026.03.10.710761

**Authors:** Praval Pratap Singh, Chandraniv Dey, Jasmeen Kaur, Sudip Chakraborty

## Abstract

We have conducted all atom molecular dynamics simulations of POPC and DPPC lipid bilayers using AMBER Lipid21 force field with eight different water models, including SPC/E, TIP3P, TIP3P-FB, TIP4P-FB, TIP4P-Ew, TIP4P/2005, TIP4P-D, and OPC, to identify the most compatible one without any modification. A number of parameters have been computed in order to understand the structure of the lipid bilayer: Area per lipid, Isothermal compressibility modulus, average Volume per lipid, electron density profile, bilayer thickness, X-ray and neutron scattering form factors, deuterium order parameter, and radial distribution function. The estimated Area per lipid, Isothermal compressibility factor, volume per lipid and bilayer thickness are highly consistent with experimental results for the SPC/E water model, indicating its suitability with the AMBER Lipid21 force field, insted of any modification. The bilayer electron density profiles of both the lipid bilayers demonstrate a little augmentation of water penetration with respect to the membrane surface for TIP4P-D water model. However, the experimental X-ray and neutron scattering form factors are aligning well with the simulated results for all studied water models, and TIP4P-D shows better for X-ray data. The deuterium order parameter for lipid acyl chains value less than 0.25 for all observed water models, depicting their disorderness for both the lipid bilayers. The lateral diffusion and reorientation autocorrelation function of the lipid molecules in both the bilayers are computed to reveal their dynamics across all water models. In comparison to other water models, the simulated trajectories predict better structure and reasonably fair dynamic properties for the SPC/E water model. The TIP4P-Ew water model reproduces the lateral diffusion co-efficient in close agreement with experiment. Reorientational dynamics for both the lipids in the bilayers for eight different water models are observed; the presence of slow and slowest time components corresponds to the lipid axial motion (wobble motion) and Twist/Splay motions. So, in view of the overall performance of the different water models with the AMBER Lipid21 all atom force field in reproducing membrane physical properties, the SPC/E water model appears to be an optimal choice.

## Introduction

Bio-membranes constitute hydrated, compositionally diverse soft-matter interfaces, exhibiting emergent material properties that result from the coupled interplay of lipid-lipid packing and lipid-water interactions across the bilayer-water boundary. ^1,2^ By regulating permeability, compartmentalization, curvature, and the free energy landscape governing protein association, membranes function as both physical barriers and active regulators of cellular function. Because interfacial hydration and headgroup electrostatics are tightly coupled to lateral lipid packing, even subtle changes in water structure or surface energetics can propagate into measurable alterations in membrane organization and mechanics. ^4,5^

Atomistic Molecular Dynamics (MD) simulation provide a practical route to resolve these coupled interactions at resolutions that complement scattering, NMR and thermodynamic observables used to characterize bilayers. ^6,7^ However, the predictive value of the membrane MD has fundamental limits according to the parameter where the forcefields must reproduce not only hydrocarbon chain packing but also headgroup conformational preferences and interfacial hydration structure in tensionless conditions. ^8,9^ Therefore, the current validation benchmarks for force fields for membranes relate to agreement with a consistent set of characteristic values for area per lipid, bilayer thickness, density/scattering profiles, order parameters and diffusion, rather than direct measures of individual headgroup values. ^10,11^ For phospholipid membranes, so called phosphatidylcholines serve as exemplary standards for which the characteristic properties of the fluid phase are experimentally well established and repeatedly brought back for validation comparisons for new force fields. ^12^ In particular, 1,2-Dipalmitoyl-sn-glycero-3-phosphocholine (DPPC) is a phosphatidylcholine phospholipid whose well characterized gel to fluid transition behavior and strong scattering data make it a challenging system for examination of phosphocholine headgroup hydration, challenging even at highly accurate simulations too. ^13,14^ The primary phase transition temperature for DPPC is about 314.15 K. ^15^ This provides an important thermodynamic benchmark that highlights subtle features of intermolecular force fields that are difficult to discern through simulation. ^13,16^ Indeed, DPPC has been a model phospholipid used extensively for bench-marking MD simulation predictions against scattering factors and NMR order parameters, allowing a clear examination of bilayer structure and dynamics. ^14,17^ To complement this fully saturated model phospholipid, 1-palmitoyl-2-oleoyl-sn-glycero-3-phosphocholine (POPC) is a monounsaturated phosphatidylcholine (16:0/18:1) model phospholipid that has been widely used for fluid bilayers nearest physiological conditions. ^18^ Having only one unsaturated tail makes POPC a good indicator of the extent to which force fields compromise conformational entropy with respect to head group packing, which may be difficult to discern when systems are fully saturated. ^19^ POPC is also a standard target in cross forcefield comparisons of phosphatidylcholine membranes, where observables such as area per lipid, hydrocarbon order parameters, and diffusion coefficients are reported side by side under matched simulation protocols. Because DPPC and POPC together span saturated and monounsaturated regimes while remaining chemically close through their shared phosphocholine headgroup, they form a compact yet informative pair for dissecting where sensitivity to interfacial water enters membrane property predictions. ^20,21^ As phosphatidylcholine headgroup geometries and hydrogen bonding patterns are fixed by interfacial water, solvent models with varying dielectric properties and interfacial energies may indirectly influence phosphatidylcholine packing and dynamics. ^22^

An often critical but underemphasized part of such a validation environment is an explicit water model. ^23^ Rigid, non-polarizable water models are parameterized against different target properties and can differ materially in surface tension, dielectric response, diffusion and hydrogen bond structure, all of which are directly relevant at lipid interfaces. ^23,24^ This becomes especially important for the zwitterionic phosphatidylcholines, where the orientation of the head group and the structure of the hydration shell affect the bilayer thickness, compressibility, and interfacial electrostatics felt by the lipid and solute molecules. ^22,25^ Consistent with this perspective, membrane simulation studies have explicitly demonstrated that changing the water model without changing the lipid parameters can shift key membrane observables and inferred transport properties. ^26,27^ Accordingly, the water models are taken as an explicit variable in benchmarking strategy and briefly summarizing the defining design features of each model, beginning with SPC/E (Extended Simple Point Charge) is an evolution of SPC that introduces an effective polarization correction missing term to better reproduce bulk energetics while maintaining a computationally efficient three site form. ^24^ SPC/E was parametrized mostly with respect to the thermodynamics of condensed phase water rather than any specific biomolecular forcefields, hence, it is often used as a general purpose baseline when separating water physics effects from solute parameterization.^2^ Transferable Intermolecular Potential with 3 Points (TIP3P) is a classic three site model introduced within a family of simple water potentials and has remained dominant in biomolecular simulation due to its low cost and deep integration into common biomolecular parameter sets. ^29^ In the CHARMM forcefield, a modified TIP3P variant (mTIP3P) was historically used during parameter development, so TIP3P family behavior is often implicitly used into many lipids and protein models. ^30,31^ On the other hand, the observed discrepancies of the TIP3P model regarding transport and dielectric phenomena led to the examination of other water models for the membranes, where the problem of permeability is fundamental to the system. TIP3P-FB ^32^ and TIP4P-FB ^32^ are Force Balance optimized refinements of three-site and four site rigid water models respectively which are designed to improve agreement with a broad set of physical properties relative to older baselines. These FB models have been discussed and reused in subsequent parametrization efforts including ion parameterization precisely because of their improved bulk property balance that can propagate into more realistic solvent solute and interfacial behaviors. ^26,27^ Importantly for membrane applications, FB water models have been tested for compatibility with established lipid force fields like CHARMM36 and shown to reproduce multiple membrane observables. ^26^ TIP4P-Ew is a four site model explicitly developed for biomolecular simulations under Ewald electrostatics, improving upon earlier TIP4P style models by targeting key bulk properties and Ewald consistent behavior ^33^ as long range electrostatics and interfacial polarization are central to membrane systems, TIP4P-Ew is frequently treated as a biomolecular oriented reference among four site models. TIP4P/2005 is a general purpose four site model parameterized to reproduce water’s temperature of maximum density and aspects of the condensed phase behavior and it is often regarded as one of the most accurate rigid models for many bulk properties. ^34^ Although TIP4P/2005 is not tied to a single biomolecular forcefield family by construction, it has been used in biomolecular contexts where improved water structure and thermodynamics are desired, including studies emphasizing balanced protein-water interactions. ^23,35^ In membrane contexts, prior work has specifically investigated TIP4P/2005 alongside other models to evaluate compatibility and its impact on interfacial dynamics near bilayers. ^26^ TIP4P-D was introduced to address deficiencies in dispersion interactions in common water models that can bias solvated biomolecules, particularly intrinsically disordered proteins, toward overly compact states. ^36^ By increasing the effective water dispersion contribution while maintaining compatibility with existing models, TIP4P-D aims to improve the balance between solute-solute and solute-water interactions. ^36^ As interfacial membranes are dispersion and electrostatics sensitive environments, TIP4P-D is a relevant candidate for testing whether dispersion corrected water shifts bilayer structural signatures through modified hydration energetics. ^26,36^ OPC (Optimal Point Charge) represents an alternative design philosophy, rather than tuning parameters primarily through conventional geometry constraints, OPC optimizes the point charge distribution to better reproduce water electrostatics and a targeted subset of bulk properties. ^37^ OPC was reported to improve agreement with a broad collection of bulk properties and to yield more accurate hydration free energies for small molecules compared with several common rigid models. ^37^ Notably for practical biomolecular workflows, OPC water model is included in the directory of AMBER forcefield, making it straightforward to deploy within AMBER family simulation pipelines. ^38^

While the lipid community has long recognized that membrane predictions depend on the lipid force field, the water model dimension is often treated as secondary or assumed transferable. ^39^ Yet the literature increasingly indicates that water model choice can influence transport, dielectric response and interfacial structure in ways that matter for membranes and for membrane proximal phenomena. ^26,27^ This motivates a benchmarking strategy that varies only the water model while keeping the lipid parameters fixed, thereby attributing differences in membrane observables specifically to the solvent representation rather than to lipid re-parameterization. ^26,27^ Therefore, quantifying solvent sensitivity is most meaningful when the lipids are held fixed within a well-validated lipid force field framework, allowing changes in observables to be attributed specifically to the water description. ^10^ Within the AMBER family, lipid forcefield development has progressed through modular frameworks toward production parameter sets designed for tensionless bilayer simulations ^40 41 42^. Lipid21 ^43^ an updated AMBER’s modular lipid forcefield enabling tensionless simulations of multiple lipid types with more complex membrane compositions and the revised headgroup torsion parameters that improve agreement with NMR order parameters as compared to the Lipid14 ^42^ force field, alongside updated hydrocarbon parameters intended to better match phase transition behavior. ^43^ The Lipid21 study further emphasizes broad structural accuracy across multiple lipid classes with standard validation observables such as areas per lipid and bilayer thickness. ^43^ Although Lipid21 shows improvements over Lipid14, the dependence of its performance on the choice of water model has not been systematically assessed across modern three-site and four-site water models using experimentally validated observables. ^8^

To address this gap, we carry out molecular dynamics simulations of POPC and DPPC bilayers using AMBER Lipid21 ^43^ parameters, with systematically varying the water models, namely SPC/E, ^24^ TIP3P,^44^ TIP3P-FB,^32^ TIP4P–FB,^32^ TIP4P-Ew,^33^ TIP4P/2005,^34^ TIP4P-D,^36^ and OPC. ^37^ We evaluate a standard, validation oriented panel of observables that connects directly to experimental constraints and established membrane forcefield benchmarking practice rather than relying on any single metric. ^45^ The structural parameters includes area per lipid, isothermal area compressibility modulus and volume per lipid, electron density profiles and thickness of bilayer. The calculated results also include observables that can be directly compared with scattering-informed membrane structures and enable estimation of the interfacial density distribution and bilayer type through the computation of X-ray and neutron scattering form factors. ^14^ The interfacial hydration is analyzed in terms of the radial distribution functions of water molecules with respect to the chemically specific sites of lipid head groups, including the phosphate and carbonyl groups. ^22^ Dynamical behavior is assessed using lateral diffusion and reorientational autocorrelation functions, linking solvent dependent interfacial physics to lipid mobility and relaxation processes that underpin membrane fiuidity. ^46^ In summary, the like-to-like design presented here allows us to estimate the uncertainty of water-model predictions in Lipid21 membrane simulations and helps to identify the water models most consistent with experimentally validated structural and dynamical properties of phosphatidylcholine bilayers.

## Methodology

### System Setup and Simulation Details

The Packmol software ^47^ is used to generate initial arrangements of the POPC and DPPC lipid bilayers, along with different water models. A total of eight different water models are used for modeling the interaction parameters of water molecules, namely: SPC/E, TIP3P, TIP3P-FB, TIP4P-FB, TIP4P-Ew, TIP4P/2005, TIP4P-D and OPC(4 point charge), to simulate POPC and DPPC lipid bilayers. Individual POPC and DPPC lipid bilayers corresponds to eight different systems, and each system includes 128 lipids with 5120 water molecules in a simulation box of dimension 7.1 nm×7.1 nm×11.6 nm for POPC and 7.1 nm×7.1 nm×10.6 nm for DPPC lipids. The above composition ensures a well-hydrated lipid bilayer structure with the hydration number of 40 and prepared exactly the same as suggested in Lipid21^43^ reference. All the sixteen systems were then subjected to Molecular Dynamics simulations using the Gromacs 2018.3 version of the parallel MD simulation package.^48,49^ The lipid forcefield used for the simulation is Amber Lipid21.^43^ The topology and parameters were generated with the help of CHARMM-GUI builder. ^50-52^ The simulation experiment on the POPC and DPPC lipid bilayers were started by eliminating the unfavorable contacts via steepest descent energy minimization techniques. The systems were then equilibrated at constant temperature (T = 303K for POPC and T=323K for DPPC). For randomization of lipid molecules, Individual NVT run was performed for a duration of 100ps, followed up with a constant temperature (T = 303K for POPC and T=323K for DPPC) and pressure (P = 1.0 atm), NPT simulation run for 60 ns during equilibration is also performed. The final equilibration of the system was continued via a production run of 500 ns long run in the NPT ensemble. The trajectories were stored for every 10ps time frame.

During the course of the simulation, the LINCS ^53^ algorithm was used in order to put constraints on H-bonds in all the lipid bilayer systems. A short range cutoff of 1.0 nm was considered for the Lennard-Jones (LJ) interaction, and the Particle Mesh Ewald (PME) method^54,55^ was employed for the long range electrostatic interactions, with cubic interpolation of the order of 4 and a fourier spacing of 0.12 nm. The real space cutoff for all simulations was 1.0 nm. PME and LJ are involved in the determination of pair-wise additive site-site contributions. Verlet cutoff scheme was employed to determine the neighbors with a cutoff of 1.0 nm with every 10 step neighbor list update. The LeapFrog integrator algorithm was used to solve Newton’s equation of motion to generate the time reversible trajectory with a time step of 2 fs. A long-range analytical dispersion correction was applied to the energy and pressure. The temperature is maintained at 303K for POPC and 323K for the DPPC lipid bilayers which are higher than their respective melting points^56,57^ using the Stochastic velocity-rescaling thermostat of relaxation time 1.0 ps. Velocity Rescale thermostat is a modified Berendsen thermostat with a stochastic term added, and the velocities are scaled by a factor lambda in order to attain the desired temperature.^58^ Parrinello-Rahman Barostat was used for Pressure Coupling for individual simulated system^59,60^ for final NPT ensemble run with a semi-isotropic coupling constant of 5.0 ps and compressibility *κ*_*xy*_ = 4.5 × 10^−5^ *bar*^−1^, and *κ*_*z*_ = 0.0. Simulated boxes adhered to the Periodic Boundary Conditions in all three dimensions (3D PBC). The structural and dynamical analysis were conducted with the help of GROMACS analysis tools ^48,49^ and SIMtoEXP software. ^61^ In this article, we have used the VMD (visual molecular dynamics) package ^62^ to produce all of the images.

## Results and Discussion

The performance of the AMBER Lipid21 force field is systematically evaluated with respect to the choice of water model using experimentally validated observables. Which is likely to control the structure and dynamics of lipid bilayers. In the following subsections, we examined the structure and dynamics of POPC and DPPC lipid bilayers using AMBER Lipid21 force field for eight different water models, namely SPC/E, TIP3P, TIP3P-FB, TIP4P-FB, TIP4P-Ew, TIP4P/2005, TIP4P-D and OPC.

### Structural Properties

#### Area per Lipid

Area per lipid (APL) is a standard structural parameter to validate the lipid bilayer simulation to make sure the simulated systems were in equilibrium. ^41^ From membrane simulations, we can effortlessly determine the Area per Lipid and whether a lipid bilayer is in the correct phase at a particular temperature. ^42^ The area per lipid can be found using experimental data. Still, as illustrated by Poger and Mark, there is often a high degree of uncertainty in experimentally derived areas per lipid values. ^45^ A wide range of APL values for a particular lipid bilayer is reported in the literature, ^63,27^ That is why performing a simulation study is essential to correctly predict the area per lipid value that should at least fall in the experimental range. The following formula applies to get the average area per lipid:

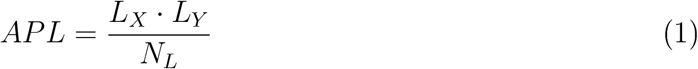

where APL, *L*_*X*_, *L*_*Y*_, *N*_*L*_ are the Average Area per Lipid, length of our simulation box in the x-direction, length of our simulation box in the y-direction, number of lipids per leaflet respectively. The simulated area per lipid (APL) values are reported in Table 1 for different water models and compared with the experimental data. For both POPC and DPPC lipid membranes, a more substantial agreement occurs between the simulated and experimental APL values for all water models. The range of 0.61-0.67 nm^2^ for each lipid indicates the liquid crystalline phase. ^64^ This shows fluidity, high lateral diffusion, rotational mobility along with disordered acyl chains of the lipid molecules. ^65^ The highest APL value for both POPC and DPPC lipid bilayer is for TIP4P-D and lowest for TIP3P-FB water models respectively. APL is also correlated with other properties like molecular packing, compressibility, bilayer thickness, and acyl chain ordering. ^66^

**Table 1:**
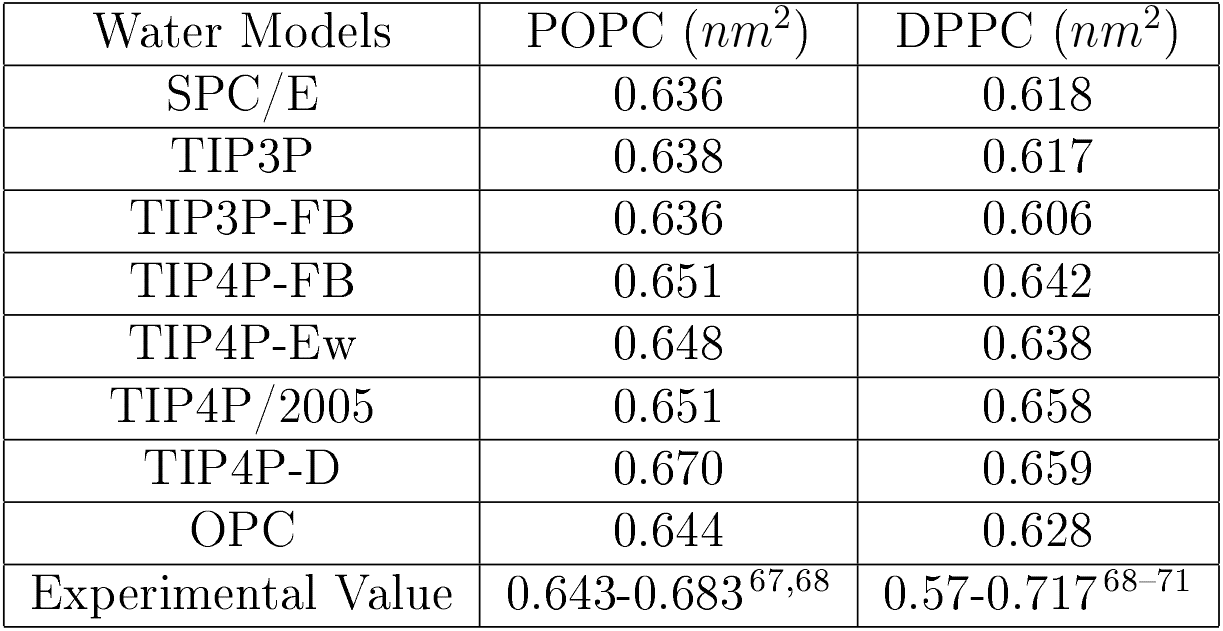
Area per Lipid Results for POPC and DPPC lipid bilayers with different water models.

#### Isothermal Area Compressibility Modulus

The isothermal area compressibility modulus describes the elastic property of membranes, and the following equation is used to calculate *K*_*A*_:

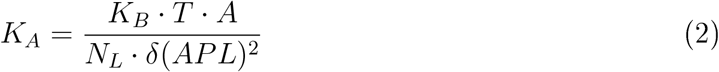

Here, *K*_*B*_ represents the Boltzmann constant, T is the average temperature, A is the average area per lipid, and *δ*(*APL*)^2^ is the variance associated with APL. The simulated average area compressibility modulus values are provided in Table 2. Which indicates that the simulated values are compatible with the experimentally reported values for the POPC and DPPC lipid bilayers, whether the SPC/E and OPC are closely consistent. On the other hand, the TIP4P-D water model for DPPC lipid bilayers shows lowest value of *K*_*A*_ compared to the other water models and depicts more fluidic and elastic behaviour. ^72^

**Table 2:** Isothermal Area Compressibility Modulus Results for POPC and DPPC lipid bilayers with different water models.

| Water Models | POPC (N/m) | DPPC (N/m) |
| --- | --- | --- |
| SPC/E | 0.307 | 0.221 |
| TIP3P | 0.239 | 0.270 |
| TIP3P-FB | 0.174 | 0.223 |
| TIP4P-FB | 0.180 | 0.240 |
| TIP4P-Ew | 0.200 | 0.171 |
| TIP4P/2005 | 0.180 | 0.170 |
| TIP4P-D | 0.344 | 0.159 |
| OPC | 0.220 | 0.220 |
| Experimental Value | 0.18-0.33<sup>73</sup> | 0.231<sup>65</sup> |

#### Volume per Lipid

The Volume occupied by lipid molecules is one of the essential parameters characterizing the lipid bilayer structure. In the simulation, the following equation calculates the Volume per lipid molecule.

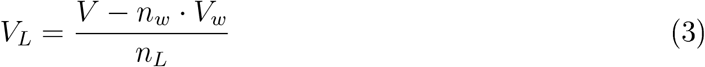

Where V represents the Volume of our simulation box, *V*_*w*_ represents Volume per water molecule, and *n*_*L*_ and *n*_*w*_ are the number of lipids (128) and water(5120) molecules, respectively. *V*_*w*_ was determined from an independent 15 ns simulation of water molecules of each water model at 303 K for the POPC lipid bilayer and 323 K for the DPPC lipid bilayer. Table 3 reports the experimental results and the average *V*_*L*_ values from the simulations. In the case of POPC and DPPC lipid bilayers, water models like SPC/E and TIP3P-FB are found in good agreement with the experimental values except TIP4P-FB, TIP4P/2005, TIP3P, TIP4P-D, OPC, and TIP4P-Ew shows little deviation from the experimental values.

**Table 3:** Volume per Lipid Results for POPC and DPPC lipid bilayers with different water models.

| Water Models | POPC ($nm^3$) | DPPC ($nm^3$) |
| --- | --- | --- |
| SPC/E | 1.23 | 1.22 |
| TIP3P | 1.20 | 1.20 |
| TIP3P-FB | 1.23 | 1.22 |
| TIP4P-FB | 1.19 | 1.18 |
| TIP4P-Ew | 1.19 | 1.18 |
| TIP4P/2005 | 1.19 | 1.18 |
| TIP4P-D | 1.20 | 1.19 |
| OPC | 1.20 | 1.18 |
| Experimental Value | 1.223-1.256<sup>67,75</sup> | 1.226-1.232<sup>65,67</sup> |

The slightly lower values of Volume per Lipid in both POPC and DPPC lipid bilayers may be due to the acyl chains of lipids being more extended. ^74^

#### Bilayer Electron Density

Electron density profiles (EDPs) provide detailed structural information of the lipid bilayer along the bilayer normal. The simulated EDPs of the POPC and the DPPC bilayers can be compared and validated with respect to the available X-ray and Neutron scattering experimental data. In Figure 2a and Figure 2b, we have presented the EDPs of the POPC and the DPPC bilayers for eight different water models. It is clear from figures that the lipid bilayer profiles are symmetric, indicating the proper equilibration during the time-scale of the simulations. The two primary peaks of the EDPs correspond to the phosphate group in the head group area of lipids, which can also be used to determine the Bilayer Thickness. Furthermore, a reduction in the curve is observed at the starting point, suggesting the existence of terminal methyl groups of the acyl chains in the lipid bilayers across all EDPs of the simulated bilayer systems. It is also observed from the figure that the water covers and penetrated up to the head group regions (i.e., upto phosphate and choline) and partially the glycerol regions (i.e., the carbonyls), but the terminal methyl groups in the acyl chains stay dehydrated, which correlates well with the earlier reports,^76,77^ and in agreement with experimental observations.^67,71,78^ Moreover the drop in the middle region of EDPs denotes the reduction in the interaction between the lipid bilayers. The drop in the curve for the POPC and DPPC lipid bilayer center is slightly more in the case of SPC/E, TIP3P and TIP3P-FB water models. Penetration of water from the membrane surface can also be predicted from this graph which shows SPC/E, TIP3P, and TIP3P-FB have displaced less water in the case of POPC and DPPC lipid bilayers, whereas water is more displaced in TIP4P-D water model for both the lipid bilayers.

### Bilayer Thickness (*D*_*HH*_)

Bilayer Thickness(*D*_*HH*_) is the distance between the maxima in the electron density profile. The bilayer thickness can be calculated as the head-to-head distance between the electrondense phosphate peaks. *D*_*HH*_ is a principal quantity used to identify the lipid bilayer’s elasticity, flexibility, packing, and orderliness. The thickness values obtained for both lipid systems POPC and DPPC lipid bilayer are compared with the experimentally reported values. The simulated *D*_*HH*_ values, with different water models, are presented in Table 4.

**Table 4:** Bilayer Thickness Results for POPC and DPPC Lipid Bilayers with Different Water Models.

| Water Models | POPC(nm) | DPPC(nm) |
| --- | --- | --- |
| SPC/E | 4.06 | 3.84 |
| TIP3P | 3.77 | 3.80 |
| TIP3P-FB | 4.06 | 3.92 |
| TIP4P-FB | 3.96 | 4.00 |
| TIP4P-Ew | 3.98 | 4.02 |
| TIP4P/2005 | 3.96 | 3.88 |
| TIP4P-D | 3.86 | 3.91 |
| OPC | 4.00 | 3.78 |
| Experimental Value | 3.70<sup>67</sup> | 3.83<sup>65</sup> |

Where, we have compared membrane thicknesses with the experimental value. For POPC lipid bilayers, the TIP3P water model is in good agreement with the experimental value, and for DPPC lipid bilayers, SPC/E is in close agreement with experiment. In contrast, the rest of the water model shows overestimations, specifically in POPC lipids. The overestimation may be because of headgroup size, which may influence the *D*_*HH*_ value, and the increasing chain length alter the thickness and cross-section area of the bilayers.

### Scattering form factors (X-ray and Neutron)

In order to determine lipid membrane structural properties, the Fourier-transformed form factors for the X-ray and Neutron data can be compared with experimental results. ^67^ The form factor is given by Eq. 4

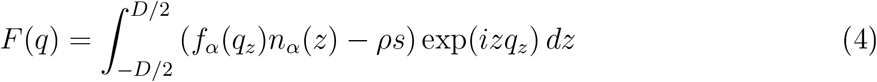

Here, *n*_*α*_(*z*) are the atomic number distributions obtained from simulations. *f*_*α*_(*q*_*z*_) is the neutron scattering length density, *ρs* is the scattering density of the solvent, and *D*_*HH*_ is the lipid bilayer thickness. In the case of X-ray scattering, *f*_*α*_(*q*_*z*_) is the atomic form factor. For centrosymmetric membranes imaginary part becomes zero, ^61^ then the Eq.4 reduces to the following form:

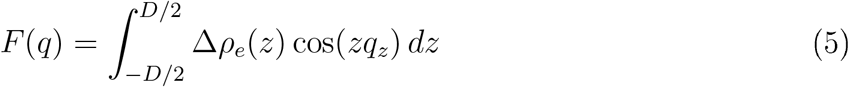

Here, Δ*ρ*_*e*_(*z*) is the scattering length density difference between the solvent and the lipid bilayer. X-ray and Neutron form factors can be determined using the SIMtoEXP software. ^61^ X-ray scattering form factors for the lipid bilayer systems are studied in different water models. In contrast, the neutron scattering form factors are studied in 100% *D*_2_*O* concentrations for POPC and DPPC lipid bilayers. Overall, the water models agree with the experimental curve for POPC and DPPC lipid bilayers. For the POPC lipid bilayer, the first maxima curve in the X-ray scattering profile indicates the structural properties of the lipid bilayer in which the TIP4P-D water model is very close to the peak with a position slightly different from the experiment, whether the SPC/E and other water model shows indifferent first peak position with respect to the experiment, the second maxima is reasonably a fair agreement with all studied water models, and the third maxima is correctly represented by the TIP4P-D water model. However, the DPPC lipid bilayer, the first maxima curve for the TIP4P-D and TIP4P/2005 water models are in good agreement with the experimental observations. The TIP4P-D model also reproduces the experimental curve for the second maxima, and the third maxima correctly represented by the TIP4P-FB, TIP4P-Ew, TIP4P/2005, TIP4P-D, and OPC water models. The X-ray form factors, especially at the minimum position near 0.27*nm*^−1^ value, indicate the bilayer thickness. The simulated and experimental form factors for different systems are shown in Figure 3a, Figure 3c, Figure 3b, and Figure 3d. The simulated Neutron form factors agreed with the experimental data for all water models.

#### Deuterium Order Parameters

The ordering of the *SN*_1_ and *SN*_2_ acyl chains for POPC and DPPC lipid bilayers can be determined using the Deuterium order parameter, which can be calculated using the following equation

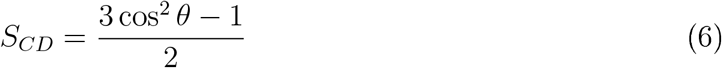

Where, *θ* is the angle between a C-D bond of the methylene group of the acyl chains and the bilayer normal. For POPC and DPPC lipid bilayers, the deuterium order parameters for *SN*_1_ and *SN*_2_ acyl chains are in good agreement with the experimental data except *SN*_2_ chain of POPC, which shows dissimilar behavior with respect to the experimental data, ^80^ as well as non-identical orientations of the *SN*_2_ chain, which are displayed in Figure 4a, Figure 4b, Figure 4c, and Figure 4d. It is important to note from the figures that for POPC and DPPC lipid bilayers, the order parameter values for all the acyl chains are lower than 0.25, representing the disordered state of aliphatic chains. ^45^ TIP4P-D water model for POPC and DPPC lipid bilayers shows more disorderness for both the acyl chains (*SN*_1_ and *SN*_2_). In Table 5 displayed the values of deuterium order parameters of headgroups. When compared to the experimental values for POPC and DPPC lipid bilayers, the deuterium order parameter for headgroups also reveals that the majority of values, such as C32(*β*), C31(*α*), and C3(*γ*_3_), for all the water models predict better results, which are labelled in Figure 1.

**Table 5:** Deuterium Headgroup results for POPC and DPPC lipid bilayers with different water models.

| Lipid Model | Water Models | C32 ($\beta$) | C31 ($\alpha$) | C3 ($\gamma_3$) | C2 ($\gamma_2$) | C1 ($\gamma_1$) |
| --- | --- | --- | --- | --- | --- | --- |
| POPC | SPC/E | 0.053 | 0.055 | 0.23 | 0.005 | 0.052 |
| POPC | TIP3P | 0.053 | 0.056 | 0.24 | 0.006 | 0.052 |
| POPC | TIP3P-FB | 0.062 | 0.048 | 0.23 | 0.0019 | 0.061 |
| POPC | TIP4P-FB | 0.046 | 0.054 | 0.24 | 0.016 | 0.062 |
| POPC | TIP4P-Ew | 0.041 | 0.056 | 0.24 | 0.004 | 0.056 |
| POPC | TIP4P/2005 | 0.039 | 0.059 | 0.25 | 0.0197 | 0.066 |
| POPC | TIP4P-D | 0.040 | 0.054 | 0.24 | 0.018 | 0.063 |
| POPC | OPC | 0.030 | 0.052 | 0.24 | 0.013 | 0.059 |
| POPC | Exp. Value | 0.050 | 0.040 | 0.21 | 0.200 | 0.130 |
| DPPC | SPC/E | 0.0512 | 0.055 | 0.24 | 0.014 | 0.061 |
| DPPC | TIP3P | 0.036 | 0.058 | 0.24 | 0.005 | 0.050 |
| DPPC | TIP3P-FB | 0.0577 | 0.052 | 0.24 | 0.022 | 0.069 |
| DPPC | TIP4P-FB | 0.0477 | 0.051 | 0.24 | 0.019 | 0.064 |
| DPPC | TIP4P-Ew | 0.0410 | 0.056 | 0.24 | 0.016 | 0.063 |
| DPPC | TIP4P/2005 | 0.039 | 0.057 | 0.24 | 0.020 | 0.068 |
| DPPC | TIP4P-D | 0.042 | 0.053 | 0.24 | 0.026 | 0.068 |
| DPPC | OPC | 0.040 | 0.052 | 0.24 | 0.017 | 0.063 |
| DPPC | Exp. Value | 0.020 | 0.050 | 0.22 | 0.210 | 0.070 |

**Figure 1:**
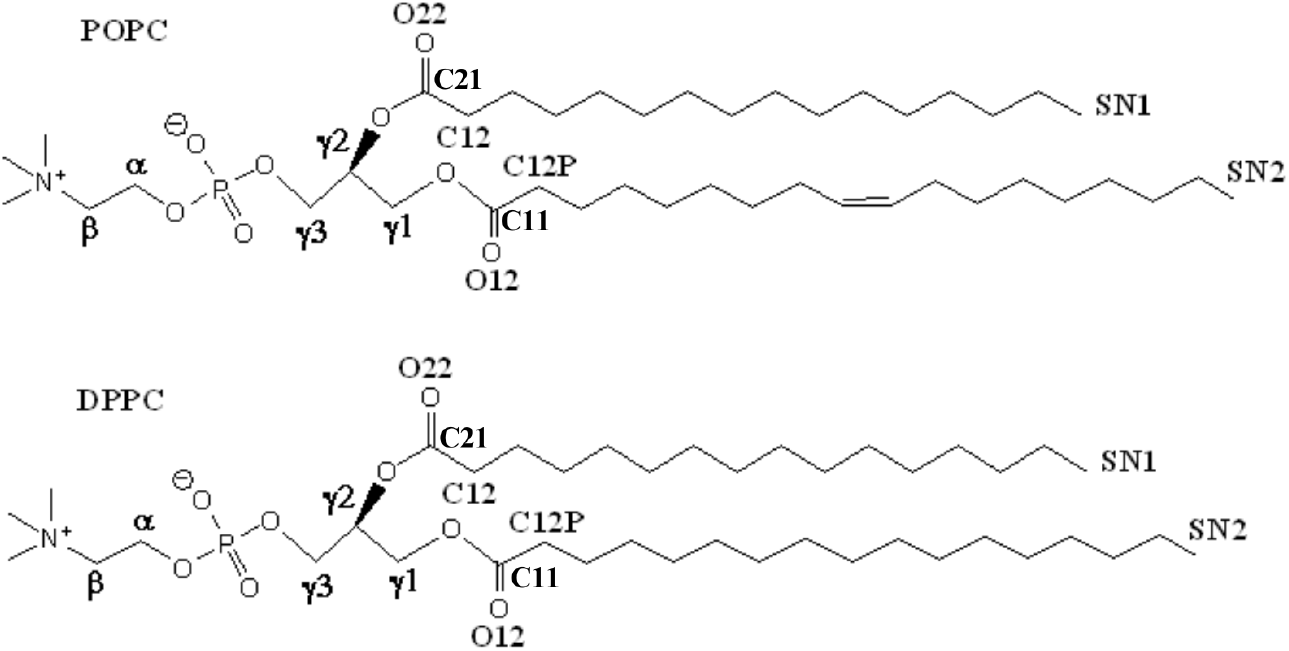
1-palmitoyl-2-oleoyl-sn-glycero-3-phosphocholine (POPC) and 1,2-Dipalmitoyl-sn-glycero-3-phosphocholine (DPPC) structures with atomic labels.

**Figure 2:**
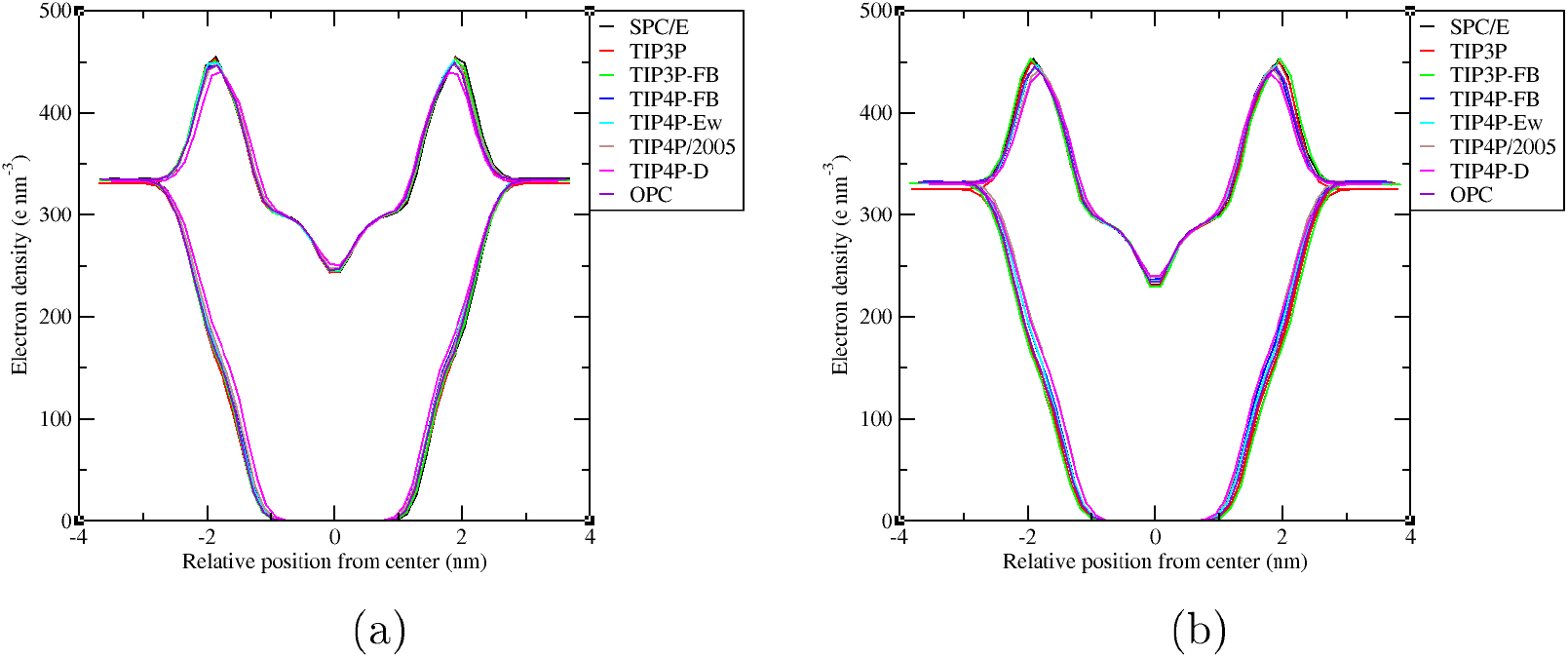
Bilayer electron density profile for (a) POPC lipid bilayers with different Water models, (b) DPPC lipid bilayers with different Water models.

**Figure 3:**
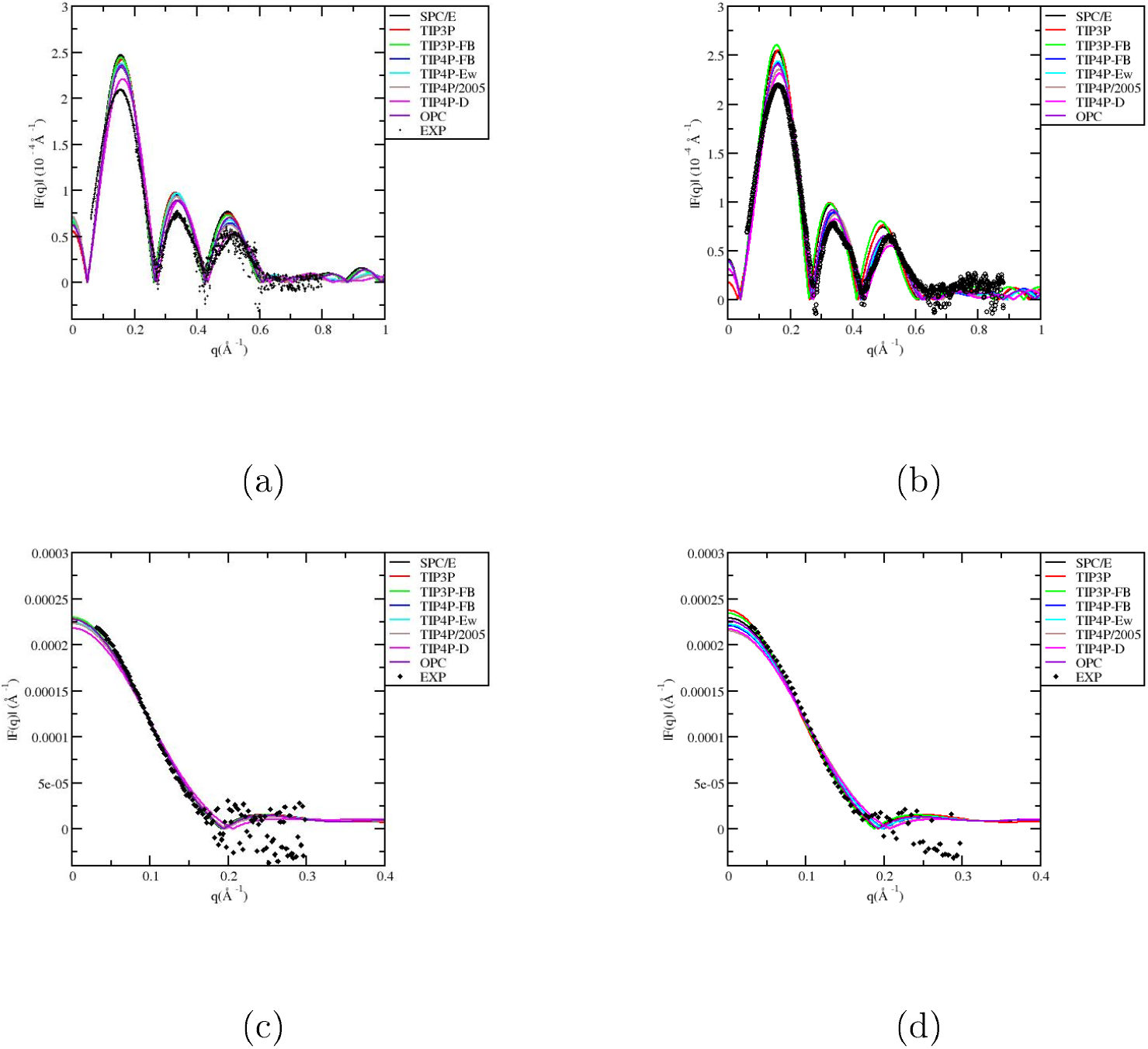
X-ray for (a) POPC lipid bilayer with different water models, (b) DPPC lipid bilayer with different water models; Neutron scattering for (c) POPC lipid bilayer with different water models, (d) DPPC lipid bilayer with different water models.

**Figure 4:**
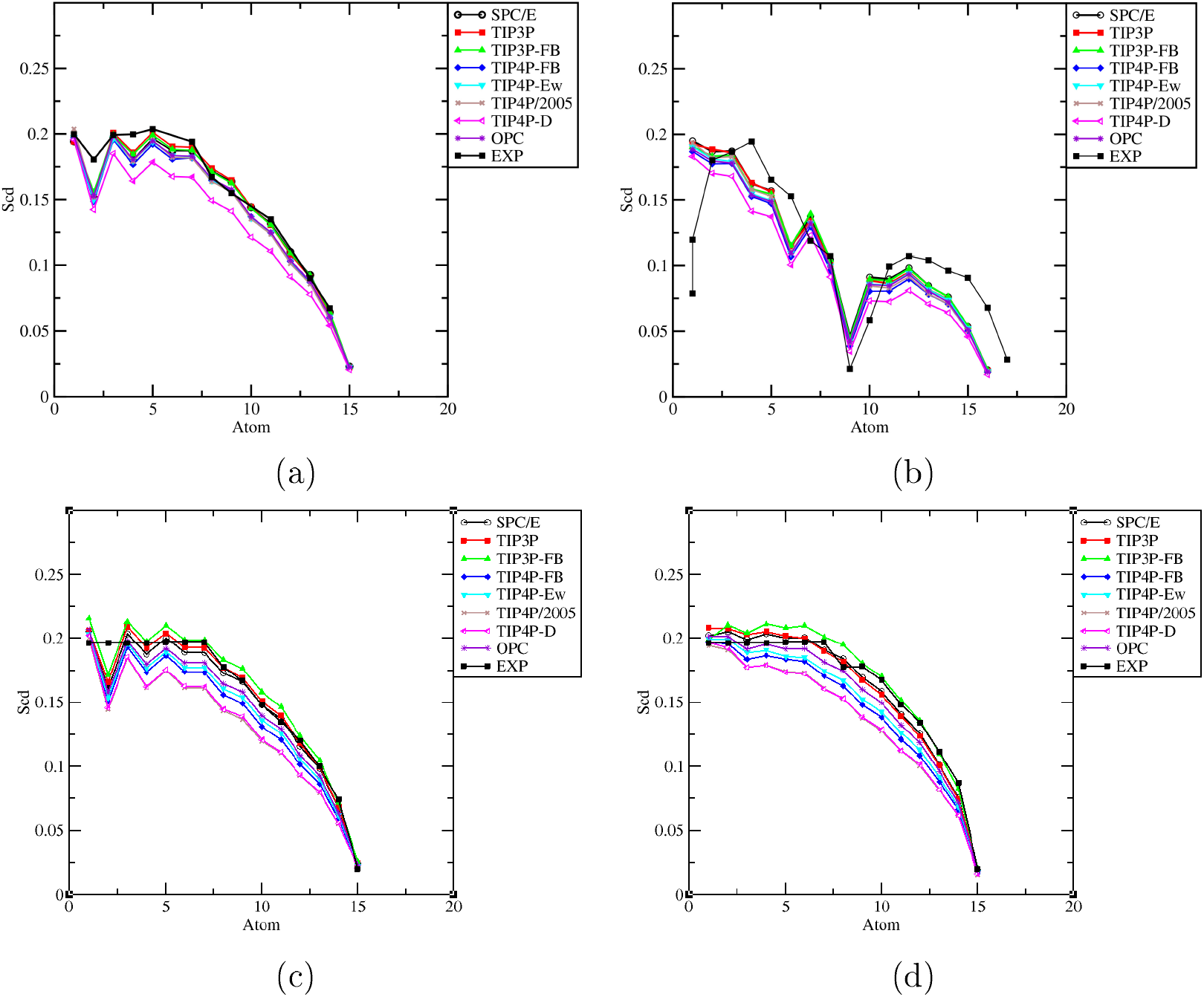
Deuterium order parameter for (a) *SN*_1_ chain of POPC lipid bilayer with different water models, (b) *SN*_2_ chain of POPC lipid bilayer with different water models, (c) *SN*_1_ chain of DPPC lipid bilayer with different water models, (d) *SN*_2_ chain of DPPC lipid bilayer with different water models.

### Radial Distribution Function

To develop understanding with the hydration structure of the lipid bilayer, we compute the radial distribution functions (g(r)) between the phosphorus (P31) atom of lipid head-group with oxygen (OW) atom of water and carbonyl oxygens of lipid (O_*C*_: O12 and O22) with oxygen (OW) atom of water for different water models, and plotted in Figure 5a, Figure 5b, Figure 5c, and Figure 5d respectively. Intriguingly, the POPC and DPPC lipid head groups have similar hydration structures, suggesting that slight modifications to the hydrophobic chains of the lipid molecules do not have much impact on the lipid hydration. The presented g(r) containing first peak intensities for both the POPC and DPPC lipids are higher for all water models, indicates the well defined first solvation shell around the lipid head groups as well as near carbonyl oxygen atoms, followed by the featureless second solvation shell. The variation in peak intensities also depicts the formation of hydration structure around lipid head group region is to a greater extent for TIP4P-D water model, and least for TIP3P, for both the lipid bilayers.

**Figure 5:**
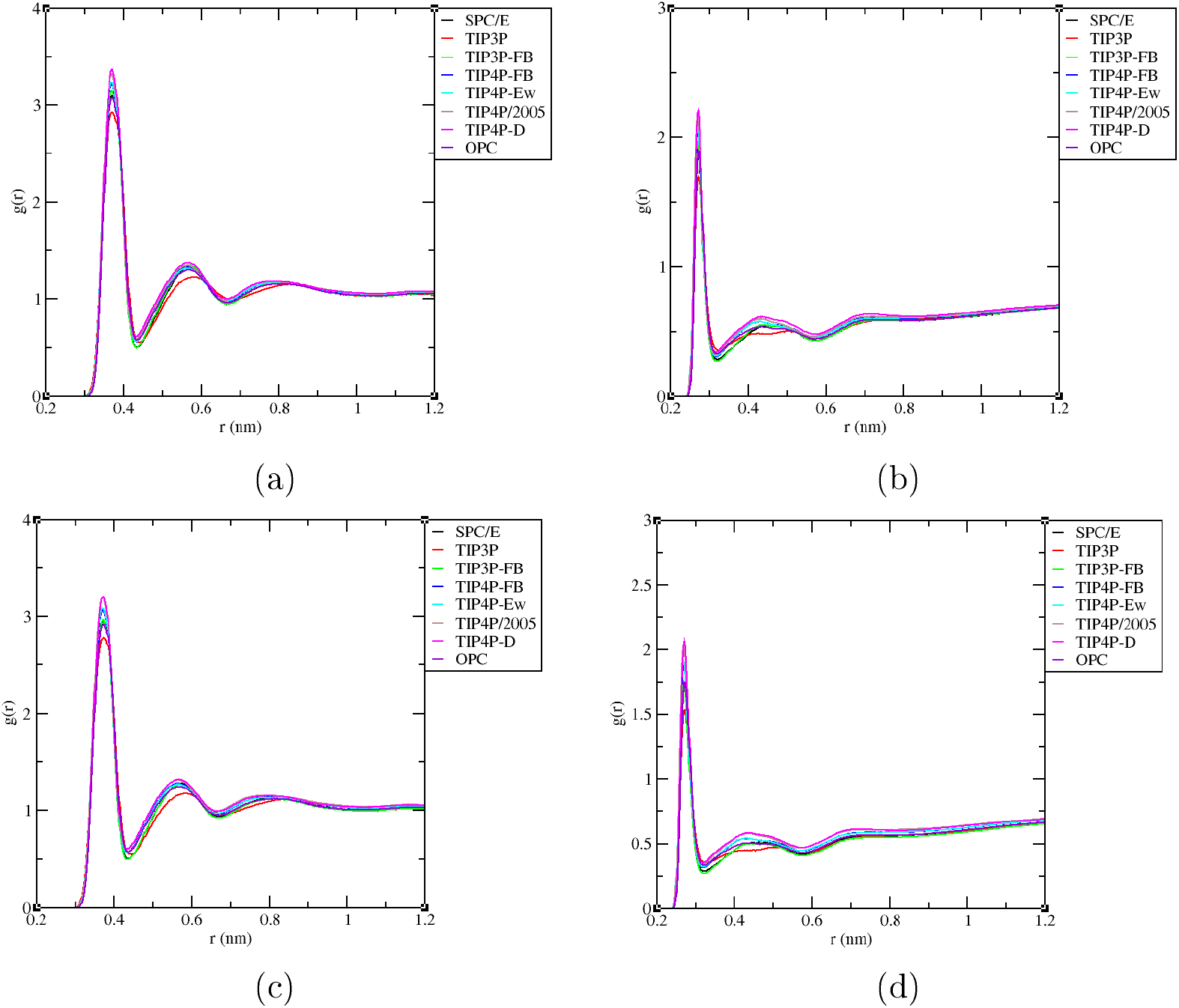
Radial distribution functions between (a) the phosphorus atom (P31) of the head-group of POPC lipid and water oxygen (OW) atom for different water models, (b) the oxygen atoms (O_*C*_: O12 - O22) of the carbonyl group for POPC lipid and water oxygen (OW) atom for different water models, (c) the phosphorus atom (P31) of the head-group of DPPC lipid and water oxygen (OW) atom for different water models, (d) the oxygen atoms (O_*C*_: O12 - O22) of the carbonyl group for DPPC lipid and water oxygen (OW) atom for different water models.

The coordination number of the first solvation shell surrounding the phosphate group, can be estimated with the integration of g(r) up to the distance corresponds to its minimum value. In the Table6, we characterizes interfacial hydration in POPC and DPPC bilayers by reporting the water coordination number (CN) and the first solvation shell width (FSSW) around two chemically distinct lipid oxygen environments, namely the phosphate associated site (P−O_*W*_) and the carbonyl oxygen site (O_*C*_−O_*W*_). The observed two consistent trends are apparent. First, the water model predominantly modulates first shell occupancy (CN) rather than the radial extent of the shell (FSSW). For the phosphate associated site, POPC exhibits P−O_*W*_ CN values spanning 2.87-3.21 and DPPC spans 2.74-3.10; TIP3P yields the lowest coordination (POPC: 2.87; DPPC: 2.74), whereas TIP4P-D yields the highest co-ordination (POPC: 3.21; DPPC: 3.10), with the remaining models are falling in between these limits. A similar dependence is observed for the carbonyl site, where POPC O _*C*_−O_*W*_ CN ranges from 0.77-0.88 and DPPC ranges from 0.73-0.86, again with TIP3P giving the lowest coordination (POPC: 0.77; DPPC: 0.73) and TIP4P-D giving the highest coordination (POPC: 0.88; DPPC: 0.86). In contrast, the FSSW values are nearly invariant across models and lipid types: for P−O_*W*_ the FSSW remains essentially fixed at 0.43 nm (with 0.44 nm only for TIP3P), and for O_*C*_−O_*W*_ it stays within 0.31-0.32 nm, indicating that differences among water models are expressed mainly as changes in the probability of first shell occupancy within an almost unchanged geometrical shell. Second, POPC is slightly more hydrated than DPPC at both sites, as reflected by consistently higher CN values for POPC across all water models specially by 0.1−0.15 for P−O_*W*_ and 0.02−0.05 for O_*C*_−O_*W*_, while the corresponding FSSW values remain comparable. Taken together, these results support an occupancy-modulation picture in which the characteristic length scale of the first hydration layer is conserved, but its population is sensitive to the choice of water model, with TIP4P-D producing the most hydrated interface and TIP3P the least hydrated interface for both the lipid bilayers.

**Table 6:** Water coordination number (CN) and first solvation shell width (FSSW) between P-*O*_*W*_ and *O*_*C*_-*O*_*W*_ atoms for POPC and DPPC lipid bilayers with different water models.

| Lipid Model | Water Model | P-$O_W$ (CN) | $O_C$-$O_W$ (CN) | P-$O_W$ (FSSW nm) | $O_C$-$O_W$ (FSSW nm) |
| --- | --- | --- | --- | --- | --- |
| POPC | SPC/E | 3.01 | 0.79 | 0.43 | 0.32 |
| POPC | TIP3P | 2.87 | 0.77 | 0.44 | 0.32 |
| POPC | TIP3P-FB | 3.04 | 0.80 | 0.43 | 0.32 |
| POPC | TIP4P-FB | 3.09 | 0.82 | 0.43 | 0.31 |
| POPC | TIP4P-Ew | 3.12 | 0.84 | 0.43 | 0.32 |
| POPC | TIP4P/2005 | 3.17 | 0.86 | 0.43 | 0.31 |
| POPC | TIP4P-D | 3.21 | 0.88 | 0.43 | 0.31 |
| POPC | OPC | 2.95 | 0.79 | 0.43 | 0.32 |
| DPPC | SPC/E | 2.90 | 0.76 | 0.43 | 0.32 |
| DPPC | TIP3P | 2.74 | 0.73 | 0.44 | 0.32 |
| DPPC | TIP3P-FB | 2.90 | 0.74 | 0.43 | 0.32 |
| DPPC | TIP4P-FB | 2.97 | 0.80 | 0.43 | 0.32 |
| DPPC | TIP4P-Ew | 3.01 | 0.81 | 0.43 | 0.32 |
| DPPC | TIP4P/2005 | 3.09 | 0.85 | 0.43 | 0.31 |
| DPPC | TIP4P-D | 3.10 | 0.86 | 0.43 | 0.31 |
| DPPC | OPC | 2.85 | 0.77 | 0.43 | 0.32 |

### Dynamical Properties of Lipid Bilayer

To investigate the influence of different water models on the translational and reorientational dynamics of POPC and DPPC lipid bilayers that simulated with respect to AMBER Lipid21^43^ all atom force field, we examined the dynamics of both the lipids for eight different water models in the following sections

#### Lateral Diffusion

The lateral diffusion of lipid molecules can be studied by monitoring the diffusion coefficient *D*_*xy*_ It helps to understand the movement of lipids within the bilayer The lateral diffusion of lipid bilayers in the membrane allows us to understand their packing and fluidity Which can be estimated from the slope of the mean square displacements (MSD) of the center of mass of the lipid vs time curve; using the well known Einstein relation. ^81^

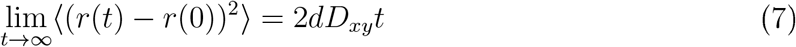

where, d is the number of dimensions (d = 2); *D*_*xy*_ is the lateral self-diffusion coefficient; and r(t) and r(0) are the center of mass positions of the lipid molecules at time t and t = 0; respectively. The averaging ⟨ … ⟩ is done over both time origins and lipid molecules.

We have calculated the mean square displacements (MSDs) of POPC and DPPC lipids along the plane of bilayer (i.e. the xy plane) for eight different water models, which are displayed in Figure 6. It is clear from the figure that the presence of fast rattling motion of the lipids ^82^ as well as the slower long time diffusion in the bilayer plane. The MSD curves were generated using 100 ns of window lengths, and diffusion coefficients were estimated as a block average based on NPT simulation runs. The calculated lateral diffusion coefficients (*D*_*xy*_) values from the MSD vs time curve, corresponding to both the bilayers with eight different water models, in the time interval 15 to 20 ns using eq 7; are listed in Table 7.

**Figure 6:**
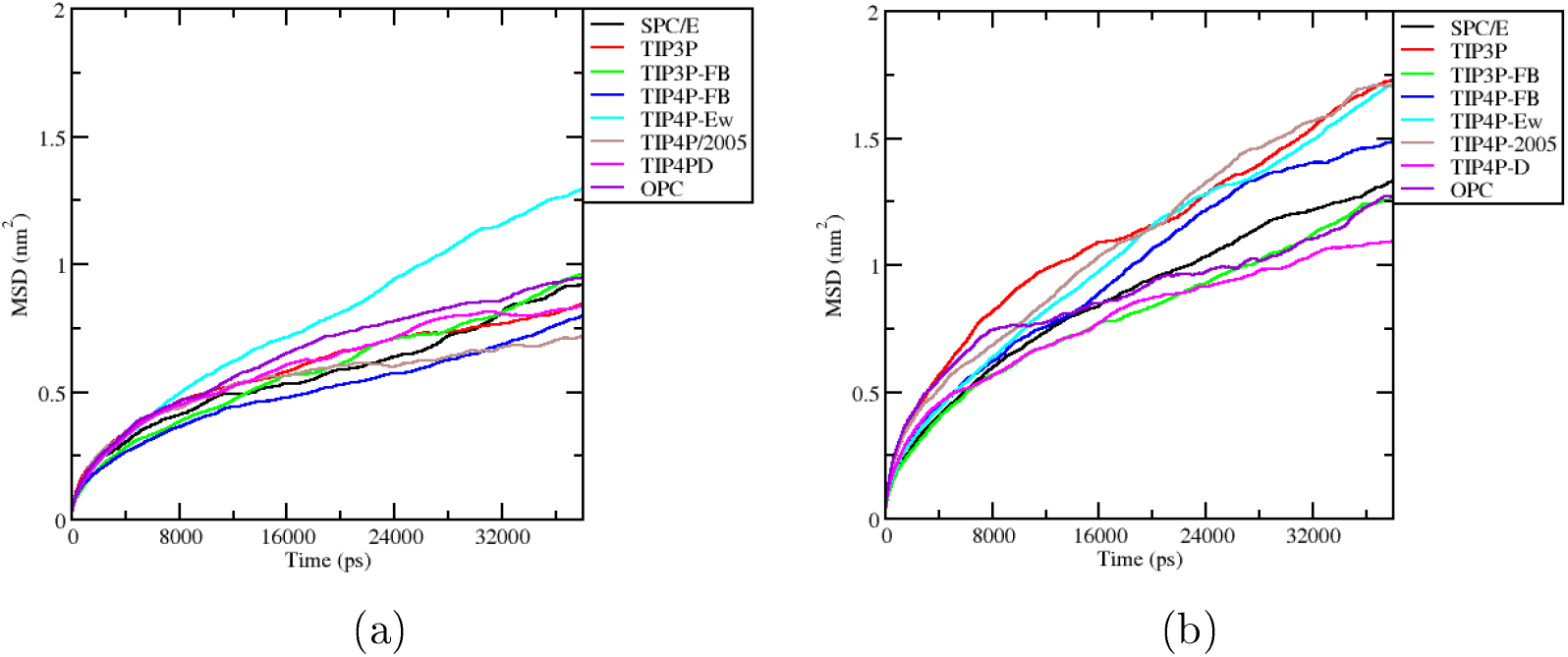
Time evolution of the in-plane (i.e. in xy plane) mean square displacements (MSDs) for (a) POPC lipid Bilayers with different water models, (b) DPPC lipid Bilayers with different water models.

**Table 7:** Lateral diffusion coefficients for POPC and DPPC lipid bilayers with different water models.

| Water Models | $D_{xy}^{POPC}$ ($10^{-8}\text{cm}^2\text{s}^{-1}$) at 303K | $D_{xy}^{DPPC}$ ($10^{-8}\text{cm}^2\text{s}^{-1}$) at 323K |
| --- | --- | --- |
| SPC/E | 3.45225 | 6.58205 |
| TIP3P | 4.84775 | 4.1634 |
| TIP3P-FB | 2.742275 | 4.378625 |
| TIP4P-FB | 3.136575 | 10.77585 |
| TIP4P-Ew | 5.89865 | 11.08395 |
| TIP4P/2005 | 2.490655 | 7.3779 |
| TIP4P-D | 2.713375 | 6.976725 |
| OPC | 5.2908 | 4.56215 |
| Experimental Value | $10.7^{96}$ | $12.5^{97}$ $15.2^{98}$ |

A variety of experimental methods have been used to study lipid lateral diffusion along the bilayer plane, including fluorescence microscopy, ^83,84^ quasielastic neutron scattering (QENS), ^85,87^ electron paramagnetic resonance spectroscopy(EPR); ^88,89^ and NMR techniques. ^90,94^ A list of values of *D*_*xy*_ for DPPC lipid span over three orders of magnitude from 0.22 ×10^−8^*cm*^2^*s*^−1^ to 460×10^−8^*cm*^2^*s*^−1^.^86^ Besides techniques, different studies using the same method also differ in their values. ^95^ Because, the experimental *D*_*xy*_ is so broad. The prediction of the lateral diffusion coefficient using a force field of lipid is difficult; due to the reason for deficiency in the force field and finite size effects. The estimated lateral diffusion coefficients; enlisted in Table 7, are found to be underestimated for TIP3P, TIP3P-FB, TIP4P/2005, TIP4P-D, and OPC water models for both the lipid bilayers. The SPC/E, and TIP4P-FB are partially; and the TIP4P-Ew water model among all studied models are shown reasonably fair agreement with earlier reported experimental results. ^96-98^

#### Reorientational Dynamics

The reorientational motions of lipid can be estimated from the reorientational autocorrelation function, which can be correlated with the relaxation time from NMR. The rotational relaxation of lipid can be influenced by the choice of water models during the course of the simuation. In this work, we have opted the C12P - C12 cross chain vector for reorientational correlation function calculations for POPC and DPPC lipid bilayers with different water models. The reorientational relaxation times are monitored by auto correlation function of 2nd order, C_2_ (t), is defined using the following equation ^99^ :

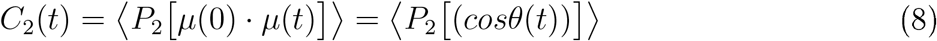

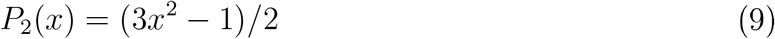

Where the order of the Legendre polynomial is 2 and *μ* represents the vector of orientation in the molecule used for the calculation. The angular brackets indicating the averaging is over both the lipid molecules and time origins. The correlation functions were fitted with multi exponentials to determine the time constants, using the three exponential decay functions ^100^ with additional exponential term (proposed in this work) to capture long time slow components.

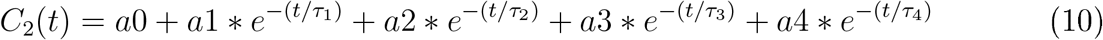

Where a0 corresponds to the average plateau value from the correlation function, a1, a2, a3, a4 are coefficients related to the four exponential decay functions. The relaxation times *τ*_1_ and *τ*_2_ corresponds to the internal lipid motions, such as gauche-trans and mid motion, *τ*_3_ represents the lipid axial motions, such as wobble motions, ^100^ and *τ*_4_ depicts the long time slow component related to the lipid bilayer Twist/Splay motions ^101^ during the time scale of the simulation.

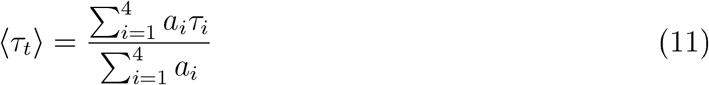

In Figures 7a and 7b, we presented the two reorientational time correlation functions, *C*_2_(*t*), of POPC and DPPC lipid in the bilayers as functions of time for different water models. The fitting parameters and the average reorientational relaxation times (⟨*τ*_*t*_ ⟩) obtained from multi-exponential (eq 10) are presented in Table 8. Which contains the presence of slow and slowest time constants irrespective of all water models, corresponds to the lipid axial motion (wobble motion^100^) and Twist /Splay motions. Our observations also indicate the presence of internal lipid motions represented by *τ*_1_ and *τ*_2_ time constants. The differential behavior of average reorientational relaxation times (⟨*τ*_*t*_ ⟩) for both the lipid bilayers with eight different water models depicts the alteration of hydrogen bonding pattern and their network relaxation times may influence the lipid dynamics. In this regard, the TIP3P water model shows faster lipid reorientational behavior for both the lipid bilayers as compared to the other water models, indicating their dynamically rapid nature and quicker network relaxations are influencing the lipid reorientational dynamics.

**Table 8:** Multiexponential fitting parameters for the reorientational correlation function of POPC and DPPC lipid in the bilayers with different water models.

| Lipid Model | Water Model | $a_1$ | $\tau_1$ (ps) | $a_2$ | $\tau_2$ (ps) | $a_3$ | $\tau_3$ (ps) | $a_4$ | $\tau_4$ (ps) | $\langle\tau_t\rangle$ (ps) |
| --- | --- | --- | --- | --- | --- | --- | --- | --- | --- | --- |
| POPC | SPC/E | 0.05 | 100.23 | 0.10 | 775.32 | 0.20 | 4198.39 | 0.48 | 28285.50 | 14474.16 |
| POPC | TIP3P | 0.08 | 114.61 | 0.13 | 741.25 | 0.27 | 3700.66 | 0.35 | 18682.50 | 7698.94 |
| POPC | TIP3P-FB | 0.05 | 112.94 | 0.06 | 720.25 | 0.20 | 3902.25 | 0.51 | 28029.60 | 15041.20 |
| POPC | TIP4P-FB | 0.06 | 100.79 | 0.09 | 750.25 | 0.23 | 3752.00 | 0.44 | 23613.00 | 11358.12 |
| POPC | TIP4P-Ew | 0.07 | 90.24 | 0.10 | 799.24 | 0.18 | 3046.92 | 0.48 | 19722.20 | 10086.93 |
| POPC | TIP4P-D | 0.06 | 95.26 | 0.09 | 723.39 | 0.23 | 3273.82 | 0.46 | 22453.70 | 11100.37 |
| POPC | TIP4P/2005 | 0.06 | 100.25 | 0.12 | 756.52 | 0.23 | 4052.02 | 0.43 | 23316.70 | 11076.23 |
| POPC | OPC | 0.07 | 120.24 | 0.08 | 790.23 | 0.25 | 4252.23 | 0.42 | 26076.90 | 12193.17 |
| DPPC | SPC/E | 0.08 | 100.25 | 0.17 | 799.35 | 0.20 | 3500.27 | 0.39 | 12693.70 | 5776.56 |
| DPPC | TIP3P | 0.13 | 113.12 | 0.19 | 730.00 | 0.33 | 3985.00 | 0.19 | 10468.80 | 3460.34 |
| DPPC | TIP3P-FB | 0.07 | 101.25 | 0.14 | 719.24 | 0.21 | 3900.25 | 0.43 | 15768.10 | 7630.36 |
| DPPC | TIP4P-FB | 0.09 | 92.95 | 0.16 | 719.26 | 0.33 | 3902.26 | 0.26 | 15764.00 | 5536.98 |
| DPPC | TIP4P-Ew | 0.11 | 97.17 | 0.19 | 799.22 | 0.19 | 3402.26 | 0.35 | 9806.81 | 4232.30 |
| DPPC | TIP4P-D | 0.09 | 101.26 | 0.18 | 740.26 | 0.32 | 3900.25 | 0.25 | 14269.10 | 4946.41 |
| DPPC | TIP4P/2005 | 0.10 | 89.62 | 0.20 | 750.26 | 0.28 | 3535.99 | 0.27 | 12136.00 | 4401.61 |
| DPPC | OPC | 0.10 | 101.24 | 0.18 | 891.38 | 0.21 | 3600.25 | 0.37 | 13483.90 | 5882.62 |

**Figure 7:**
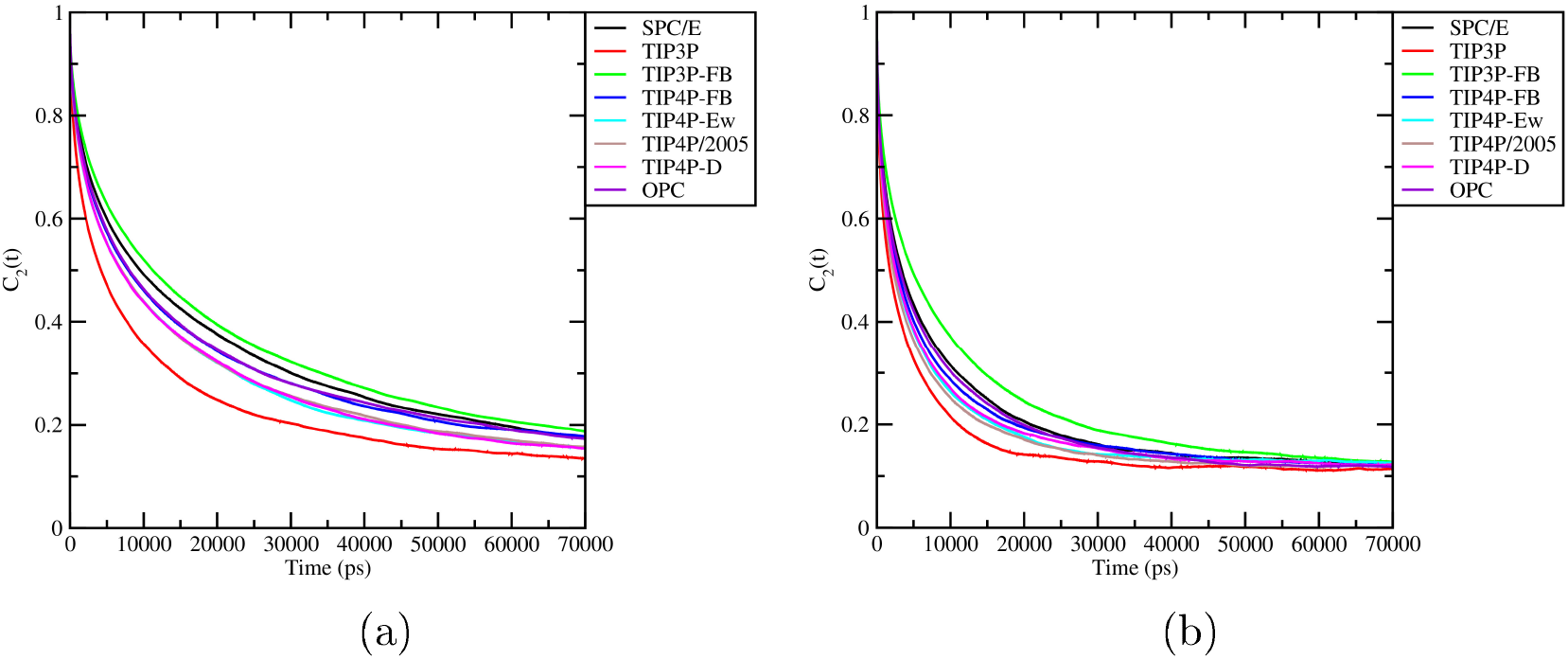
Reorientational time correlation functions of the C12P – C12 cross chain vector for (a) POPC lipid in the bilayer with different water models, (b) DPPC lipid in the bilayer with different water models.

## Conclusion

In this article, we have carried out molecular dynamics simulations of POPC and DPPC lipid bilayers using AMBER Lipid21 ^43^ all atom force field with eight different water models, namely SPC/E, ^24^ TIP3P,^44^ TIP3P-FB,^32^ TIP4P-FB,^32^ TIP4P-Ew,^33^ TIP4P/2005, ^34^ TIP4P-D,^36^ and OPC. ^37^ A methodical analysis of the lipid bilayer’s structure and dynamics has been conducted. To comprehend the structure of the lipid bilayer, various parameters have been calculated, including area per lipid, isothermal compressibility modulus, average volume per lipid, electron density profile, bilayer thickness, X-ray and neutron scattering, deuterium order parameters of the hydrocarbon chains and head groups, and radial distribution function. To understand the dynamics of the lipid bilayer, the lipid molecules’ lateral diffusion and reorientation autocorrelation function are computed. The simulations predict better structure and reasonably fair dynamic properties for the SPC/E water model as compared to the other studied water models. In comparison to the SPC/E, the TIP4P-Ew water model predicts the lateral diffusion co-efficient in close agreement with experimental results. The simulated APL parameters for both the lipid bilayers with respect to all water models are in good agreement with the experimental value. Among all studied water models, the SPC/E shows a reasonably good agreement with the experimental observations for isothermal compressibility factor, volume per lipid and bilayer thickness, and some of the water models shows partial alignment. The bilayer electron density profile represents an increased penetration of water with respect to the membrane surface as well as augmented interaction between the lipid headgroups and the water for TIP4P-D as compared to the other studied water models. The scattering form factors like X-ray and neutron scattering experimental data’s are aligning well with simulated data for all observed water models, and TIP4P-D water model produces better agreement with X-ray experiment. We observed the deuterium order parameter for acyl chains (S_*CD*_) value less than 0.25 for all studied water models, depicts the disorderness of *SN*_1_ and *SN*_2_ acyl chains for both the lipids, where TIP4P-D data reflects more such disorderness. Which can be correlated with more elasticity and liquid crystalline behaviour. The radial distribution functions for 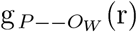 and 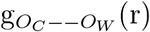 shows the first peak intensity is maximum for TIP4P-D water model, whereas the TIP3P featured the less intensity. Interestingly, the higher co-ordination number of water is observed for TIP4P-D water model for both the lipid bilayers, depicts the more availability of water around the lipid headgroup regions. Which may signifies the extended nature of molecular interaction for TIP4P-D water model, supporting the nature of bilayer electron density profile. The calculated lateral diffusion co-efficients for POPC and DPPC lipids along the plane of bilayer (i.e. the xy plane) for eight different water models (Table 7) are found to be underestimated for TIP3P, TIP3P-FB, TIP4P/2005, TIP4P-D, and OPC water models. The SPC/E, and TIP4P-FB are partially, and the TIP4P-Ew water model among all studied models are shown reasonably fair agreement with the experimental results. ^96-98^ Reorientational dynamics of both the lipids in the bilayers for different water models are observed to be faster for TIP3P water model, signifying their dynamically rapid behavior and quicker network relaxations are governing the faster lipid reorientational dynamics. So, the above observation indicates the SPC/E water model makes a balance between structure and dynamic variables during their predictions with respect to experiment.

## Acknowledgements

This work was supported in part by generous grants from the University Grants Commission (UGC)-BSR Start-up grant (F.30-432/2018 (BSR)), the Anusandhan National Research Foundation (ANRF) under the Partnerships for Accelerated Innovation and Research (PAIR) programme under sanction order No. ANRF/PAIR/2025/000029/PAIR dated 20 September, 2025 to Central University of Punjab, the Research Seed Money grant from the Central University of Punjab, and the Department of Science and Technology (DST) under the Fast Track Scheme for Young Scientists (SB/FT/CS-158/2013), Government of India. P.P.S. thanks CSIR for providing a scholarship, and C.D. thanks Central University of Punjab for providing a fellowship. J.K. thanks Central University of Punjab, for support of her Master’s project. S.C. is thankful to Vice Chancellor, Central University of Punjab, for encouraging the research and providing the necessary facilities.

